# Effects of CO_2_ on the growth and metabolism of *Akkermansia muciniphila*

**DOI:** 10.64898/2026.10.01.756130

**Authors:** M. Sakurai, S. Ogiwara, R. Osawa, J. Ying, T. Satoh

**Affiliations:** Department of Molecular Microbiology, Faculty of Life Sciences, Tokyo University of Agriculture, 1-1-1 Sakuragaoka, Setagaya-ku, Tokyo 156-8502, Japan; Department of Bioscience, Faculty of Life Sciences, Tokyo University of Agriculture, 1-1-1 Sakuragaoka, Setagaya-ku, Tokyo 156-8502, Japan

**Keywords:** *Akkermansia muciniphila*, CO_2_, metabolome

## Abstract

The human gut is home to more than 100 trillion bacteria, where various environmental factors, such as nutrients and gases, exist and influence them. The levels of fermentation gases in the gut, including CO_2_, fluctuate depending on the host’s diet. However, how CO_2_, an environmental factor, affects the physiological states of gut anaerobes remains poorly understood, but CO_2_ is often utilized in their cultivation. *Akkermansia muciniphila* (*A. muciniphila*) is widely studied as a beneficial human gut microbe. However, the effects of CO_2_ on the physiological state of *A. muciniphila* are not fully understood. In this study, we focused on CO_2_ as an environmental factor and aimed to clarify how CO_2_ affects the physiological state of *A. muciniphila*. Culture experiments showed that compared with the absence of CO_2_, the presence of CO_2_ significantly increased the growth of *A. muciniphila*. The results of the metabolomic analysis revealed distinct metabolic profiles in the presence or absence of CO_2_. The tracing analysis using stable ^13^C-labelled CO_2_ demonstrated that CO_2_ was incorporated into various metabolites in the TCA cycle, the de novo purine/pyrimidine synthesis pathways, and other pathways. In addition, the activity of pyruvate carboxylase, which catalyses an anaplerotic reaction involving CO_2_ fixation to supply oxaloacetate (OAA) to the TCA cycle, was significantly higher in *A. muciniphila* cells grown in the presence of CO_2_. Moreover, supplementation with OAA increased the growth of *A. muciniphila* in a dose-dependent manner, mirroring the effects of CO_2_ supplementation. These findings suggest that *A. muciniphila* utilizes CO_2_ for an anaplerotic reaction in the TCA cycle and other pathways involving CO_2_ incorporation, facilitating metabolic flux to promote its robust growth.

## 1. Introduction

The human intestine is inhabited by more than 100 trillion bacterial cells, known as the gut microbiota, that play a critical role in maintaining human health. The composition of the human gut microbiota is influenced by the host diet, lifestyle, genetic background, and various other factors (Ren *et al*., 2023). Among these factors, diet is known to have the most significant effect on the gut microbiota (Ross *et al*., 2024), and numerous studies have reported that a high-fat diet alters the composition and diversity of the microbiota (Murphy *et al*., 2015). In addition, intestinal gases are affected by the host diet, a high-fibre diet increases the amount of fermentative gases (CO_2_, H_2_ and CH_4_) (Modesto *et al*., 2022) and mimicking a high-fat diet decreases gas production and its production rate in human faecal culture (Li *et al*., 2022). Gases are among the human intestinal environmental factors and include N□, O□, CH_4_, H□, CO□ and other gases (Naito *et al*., 2018). Intestinal gases are supplied by host swallowing, bacterial metabolism, and diffusion from the blood to the lumen (Levitt and Bond, 1970). Their overproduction may be associated with various diseases and physiological states (Kalantar-Zadeh *et al*., 2019), such as irritable bowel syndrome (IBS) (Naito *et al*., 2018) (Major *et al*., 2017) and small intestinal bacterial overgrowth (SIBO) (Shin, 2014). Conversely, CO□ insufflation during colonoscopy is beneficial for maintaining microbiota homeostasis in patients with dysbiosis and for mitigating minor complications after colonoscopy by promoting the growth of probiotic anaerobes (Yang *et al*., 2022).

CO_2_ is present at low levels in the atmosphere (approx. 0.04% V/V), but it is present at high levels within cells owing to their respiration and plays an important role in many physiological responses across all domains of life (Tresguerres *et al*., 2010). In the gut, CO□ is one of the primary gases (approx. 10% V/V) produced during epithelial respiration and bacterial metabolism in the intestine, and its levels vary with dietary factors such as fibre intake (Modesto *et al*., 2022). CO_2_ plays a crucial role in the growth of many anaerobic bacteria, such as Lactobacilli (Valley and Rettger, 1927), *Streptococcus* (Repaske *et al*., 1974), and *Bifidobacterium* (Kawasaki *et al*., 2007) (Ninomiya *et al*., 2009), and it may promote their growth. Thus, CO_2_ is widely used as an anaerobic gas in anaerobic culture. On the other hand, some *Clostridium* strains were unaffected even when CO_2_ was completely removed by the addition of NaOH (Reilly, 1980). Thus, the effects of CO_2_ on anaerobes are not always the same and may vary depending on the bacterial species, culture conditions and medium. However, the effects of CO_2_ on various gut anaerobes and the detailed mechanisms underlying their metabolism when bacterial growth is stimulated by CO_2_ remain unclear. Understanding the interactions between CO_2_ and individual bacteria could provide new insights into gut microbiota-related host health and aid in its management.

*Akkermansia muciniphila* is a gram-negative bacterium and obligate gut anaerobe that can utilize mucin as the sole carbon and nitrogen source (Derrien *et al*., 2004), and its abundance is negatively correlated with inflammatory bowel disease (Png *et al*., 2010), diabetes (Zhang *et al*., 2013), and obesity (Karlsson *et al*., 2012). A previous study has shown that CO_2_ increases *A. muciniphila* growth and exhibits antiobesity efficacy in high-fat diet-fed mice (Wang *et al*., 2025). However, the effects of CO_2_ on the physiological state of *A. muciniphila* are not fully understood.

In this study, we performed culture experiments and a metabolomic analysis to elucidate the effects of CO_2_ on the physiological state, including the metabolomic profiles, of *A. muciniphila*. Furthermore, metabolic flux was analysed using stable ^13^C-labelled CO_2_ (^13^CO_2_) to determine in which metabolic pathway CO_2_ was utilized in *A. muciniphila*.

## 2. Materials and methods

### Bacterial strain and culture conditions

*A. muciniphila* DSM 22959^T^ was cultured in Gifu Anaerobic Medium (AccuDia™ GAM Broth; SHIMADZU, Kyoto, Japan). The medium was adjusted to a pH of 7.1 with 10 N NaOH prior to boiling. The medium was sparged with N_2_ gas, and 28 mL of anoxic medium was transferred into 120 mL serum bottles. The bottles were sealed with butyl rubber tube stoppers and autoclaved. Before inoculation, filter-sterilized *N*-acetyl-D-glucosamine (GlcNAc) (FUJIFILM Wako Pure Chemical Corporation, Osaka, Japan) was added to a final concentration of 25 mM. Filter-sterilized CO_2_ gas was injected into the headspace at final concentrations of 1, 3, 5, 10, and 20% (v/v) by withdrawing an equivalent volume of N_2_ gas to achieve various CO_2_ levels. After inoculation, the bottle was placed horizontally, and *A. muciniphila* was incubated for 24 h at 37□°C with orbital shaking at 45 rpm. The bacterial growth (OD_660_) and pH were monitored during culture.

Filter-sterilized OAA was supplemented into GAM medium (containing 25 mM GlcNAc) at final concentrations of 1, 2, 5, and 10 mM to investigate the effects of OAA. Cultivation was performed under 100% N_2_ anaerobic conditions.

### Sample preparation for the metabolomic analysis

*A. muciniphila* cells grown in GAM containing 0, 5% ^12^CO_2_ or 5% ^13^CO_2_ were harvested at the early (CO_2_ (−); 13–16 h, CO_2_ (+); 13–14 h), middle (CO_2_ (−); 17–20 h, CO_2_ (+); 16–17 h), and late (CO_2_ (−); 20–24 h, CO_2_ (+); 19–20 h) logarithmic phases. ^13^CO□ was purchased from Shoko Science Co., Ltd. (Kanagawa, Japan) (^13^C, 99.3 atom%). The culture bottles were rapidly cooled in an ice-water bath, and the cells were immediately harvested using a vacuum system and a polycarbonate membrane filter (0.2 µm pore size, 47 mm diameter; Merck KGaA, Darmstadt, Germany), with the total biomass normalized to an OD_660_ × volume (mL) = 10. The cells were then washed with ice-cold phosphate-buffered saline (PBS) on the filter membrane to remove the compounds present in the medium, and the membrane with the washed cells was placed into ice-cold methanol to quench the intracellular metabolites. The cells were detached from the membrane by sonication for 30 s and incubated for at least 10 min on ice. The samples were stored at -80 °C until further processing. These preparations were performed in three biological replicates (n=3). Metabolites were extracted from methanol-quenched cells via the Bligh–Dyer method with some modifications. Briefly, the quenched cell suspension (in methanol) was mixed with chloroform and distilled water at a ratio of 2.5:1:1 (methanol/chloroform/water, v/v/v). A procedural blank was prepared by replacing the sample with pure methanol. The mixture was incubated for 30 min at 37 °C with shaking at 1,200 rpm. Following the addition of 500 µL of distilled water, the mixture was centrifuged at 15,000 × *g* for 15 min at 4 °C to induce phase separation. The upper hydrophilic phase was collected, and the organic solvent was removed using a vacuum centrifuge, followed by lyophilization. The lyophilized metabolite samples were resuspended in 75 µL of 60% (v/v) methanol. All the samples were filtered through Ultra free-MC-PLHCC Centrifugal Filter Units (Millipore, Billerica, MA, USA) before they were subjected to LC□MS/MS. A 3.5 µL aliquot from each fully processed sample was pooled and used as a quality control (QC).

### Metabolomic analysis

The untargeted metabolomic analysis and stable isotopic tracing analysis were performed using an UltiMate 3000 Rapid Separation LC system coupled to a Q Exactive™ Focus mass spectrometer (Thermo Fisher Scientific Inc.). A ZIC-pHILIC column (2 × 150 mm, 5 µm; Merck, KGaA, Darmstadt, Germany) was used for chromatographic separation. The mobile phase was 20 mM ammonium bicarbonate (pH 9.6), labelled solvent A, and 100% acetonitrile, labelled solvent B. The following chromatographic conditions were used: injection volume, 3 µL; flow rate, 200 µL/min; and column temperature, 25 °C. Gradient elution was programmed as follows: 0–18 min, 95–60% B; 18–24 min, 60–10% B; 24–30 min, 10% B; and 30–45 min, 95% B as a re-equilibration step for the next sample.

Mass spectrometry was conducted using a heated electrospray ionization (HESI) source in polarity-switching ion mode with the following parameters: sheath gas flow rate, 10; aux gas flow rate, 0; spray voltage, 4.00 kV; capillary temperature, 320 °C; S-lens RF level, 50.0; and aux gas heater temperature, 350 °C. For full MS scans (MS1), the data were acquired at a resolution of 70,000, a scan range of *m/z* 70–1,000, and an automatic gain control (AGC) target of 1 × 10^6^. Data-dependent MS/MS (dd-MS2) spectra were acquired at a resolution of 17,500 with an isolation window of 2.0 *m/z*. Stepped normalized collision energies (NCEs) were set at 10, 15, and 20, with an AGC target of 5 × 10^4^. All the samples were analysed in three biological replicates (n=3), and all the acquired raw data were processed for baseline correction, alignment, and compound annotation using Compound Discoverer 3.3 (CD3.3) (Thermo Fisher Scientific, Inc.).

### Processing of the metabolomic data

For the untargeted metabolomic analysis, the data from 0 and 5% ^12^CO_2_ samples were used, and the template workflow “Untargeted Metabolomics with Statistics Detect Unknown with ID using Online Databases and mzLogic” in CD3.3 was used with some modifications. Compounds were detected with a mass tolerance of 5 ppm, a minimum peak intensity of 10,000, and a signal-to-noise (*S/N*) threshold of 1.5. The following adducts were considered for compound assembly: [*M* + *H*]^+^, [*M* + *H* + *MeOH*]^+^, [*M* + *NH*_*4*_]^+^, and [*M* -*H*]^-^. Compounds were grouped with a mass tolerance of 5 ppm and a retention time (RT) tolerance of 0.2 min, requiring detection in at least three files with a peak rating ≥ 3. Metabolite annotation was performed with 10 ppm tolerance by referencing multiple databases: mzCloud, mzVault, BioCyc, HMDB, KEGG, and Metabolika. For mzVault, the “Bamba lab 598 polar metabolites stepped NCE 10 30 45 library” was used, and reference spectral data for succinic acid, FAD, and CoA were newly registered in the mzVault library to ensure precise metabolite matching. For each sample, compounds with an “Annot. DeltaMass [ppm]” ≥10 ppm, unannotated compounds, and compounds with hits in neither mzCloud nor mzVault were excluded from the final dataset for subsequent analysis. Principal component analysis (PCA) was performed, and a heatmap of the hierarchical clustering analysis and a volcano plot were generated using Metaboanalyst 6.0.

For stable isotopic tracing analysis, the template workflow “Stable Isotope Labelling with Metabolika Pathways and ID using Online Databases” in CD3.3 was applied with some modifications that were the same as those in the untargeted metabolomic analysis (change to “True” in the “Match Ion Activation type” and “Match Ionization Method” nodes). The data from the 5% ^12^CO_2_ and 5% ^13^CO_2_ samples were included in this analysis as “unlabelled” and “labelled” samples, respectively. Using both “unlabelled” and “labelled” data, the incorporation ratio and the number of isotopic ^13^C atoms were calculated.

### Assessment of pyruvate carboxylase (PCx) activity

*A. muciniphila* was cultivated in 60□mL of GAM with 25 mM GlcNAc in the presence of 0% or 5% CO□ at 37 °C. Bacterial cells were harvested in the stationary phase by centrifugation (15,000□×□*g*, 15□min, 4 °C), washed with PBS, and resuspended in 10 mM Tris-HCl buffer (pH 7.8) to a final volume of 3 mL. The cells were then disrupted using a Multibeads Shocker (Yasui Kikai Corporation., Osaka, Japan) at 2,700 rpm and 4 °C for 5 cycles of 60 s on/90 s off, and cell-free extracts (CFEs) were obtained by collecting the supernatant after centrifugation (15,000□×□*g*, 15□min, 4 °C). The protein concentration in each sample was determined using the Bio-Rad Protein Assay Dye Reagent (Bio-Rad Laboratories, Hercules, CA, USA). All procedures were performed on ice to avoid the unintentional inactivation of enzyme activity. Using CFEs, pyruvate carboxylase (PCx) activity was measured spectrophotometrically (U-3900H, Hitachi High-Tech Corporation). The reaction was performed at 37 °C in a mixture containing 10 mM Tris-HCl (pH 7.8), 2.5 mM pyruvate, 25 mM NaHCO□, 0.2 mM NADH, 2 mM ATP, 150 mM KCl, 0.1 mM acetyl-CoA, 5 mM MgSO□, and 10 U of malate dehydrogenase (from Yeast; FUJIFILM Wako Pure Chemical Corporation, Osaka, Japan). The reaction was initiated by adding CFEs.

Endogenous NADH oxidation activity was also measured by performing a spectrophotometric assay in 10 mM Tris-HCl (pH 7.8) solely containing 0.2 mM NADH. The PCx activity [mU] was calculated based on the decrease in micromoles of NADH (*A*_340_) per minute after the endogenous NADH oxidation activity was deducted. The specific activity was normalized to the protein concentration and is reported as [mU/mg].

### Statistical analysis

All the statistical analyses were performed using IBM SPSS Statistics (version 25; IBM Corp., Armonk, NY, USA). The data corresponding to the culture experiments and enzymatic activity are presented as the means ± standard deviations (SDs) of at least three independent biological replicates. For comparisons among multiple groups, one-way analysis of variance (ANOVA), followed by Tukey’s post hoc test, was employed. Comparisons between two groups were conducted using a two-tailed unpaired Student’s t test. In all cases, a *p* value□< 0.05 was considered to indicate statistical significance.

## 3. Results

### CO_2_ increased the growth of *A. muciniphila*

The effect of CO_2_ on the growth of *A. muciniphila* was examined by growing the cells in GAM containing different amounts of CO_2_. Compared with 0% CO_2_, the presence of CO_2_ significantly increased *A. muciniphila* cell growth; growth stimulation was evident even at 1% CO_2_, with maximum growth observed at 5% CO_2_ or higher (Figure 1A). The pH also decreased in association with growth in the presence of CO_2_ at different concentrations (Figure 1B). *A. muciniphila* cells grew well in the presence of CO_2_ in CP medium, similar to their growth in GAM (Figure S1A and B). Furthermore, the cells grown in CP medium with 0% CO_2_ grew slower than those grown in the presence of CO_2_ but eventually reached a similar level of growth. Since *A. muciniphila* could grow in GAM even under 0% CO_2_ conditions and showed a significant difference in growth between the absence and presence of CO_2_, GAM was selected as the basal medium for subsequent experiments.

**Figure 1.**
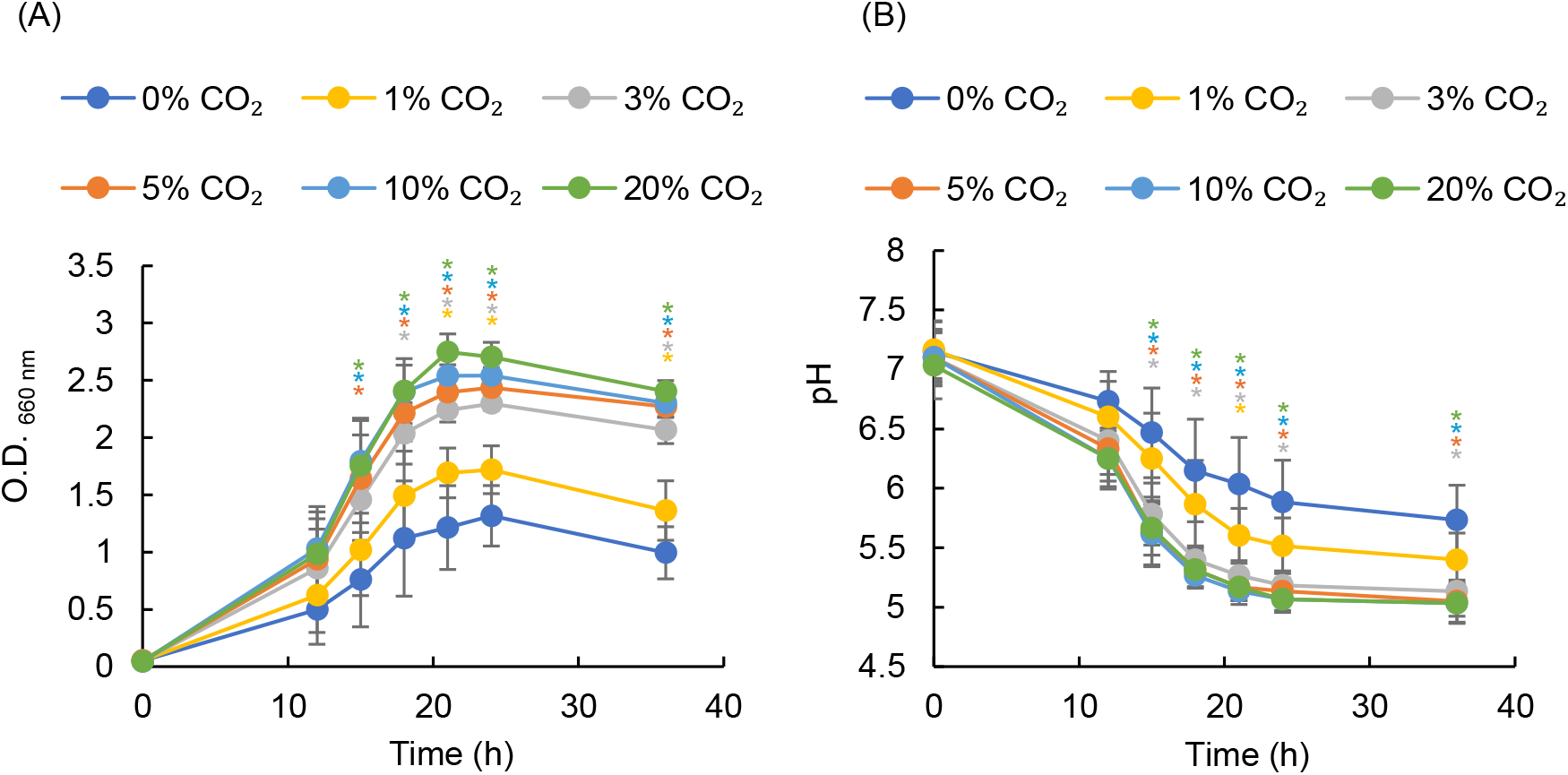
Effects of CO_2_ on the growth of *A. muciniphila*. Growth of *A. muciniphila* (A) and changes in pH (B) in the presence of 0%, 1%, 3%, 5%, 10% and 20% CO_2_. The data are presented as the means ± SDs of six biological replicates. Statistical significance was determined by Tukey’s HSD test (\**p* < 0.05, vs. 0% CO_2_).

[Figure 1. Effects of CO_2_ on the growth of *A. muciniphila*.]

### CO_2_ altered the metabolic profile of *A. muciniphila*

We investigated the effects of CO_2_ on the intracellular metabolism of *A. muciniphila* by performing a metabolomic analysis using cells grown under 0% and 5% CO_2_ conditions, designated as CO_2_ (−) and CO_2_ (+), respectively. The fraction of hydrophilic metabolites was extracted and subjected to LC□MS/MS. A total of 2,073 metabolic features were detected (Table S1), and 295 compounds annotated with high match scores were subsequently analysed (Table S2). Principal component analysis (PCA) was performed to distinguish the global metabolic profiles among the samples (Figure 2A). The distance on the score plot reflects the degree of the differences among individual samples. The results indicated a trend towards metabolic clustering between CO_2_ (−) and CO_2_ (+). A heatmap of the results of the hierarchical clustering analysis indicated the relative levels of these metabolites (Figure 2B). Volcano plots were constructed to compare metabolite levels between CO_2_ (−) and CO_2_ (+) conditions in each growth phase (Figure 2C–E). Metabolites corresponding to a raw *p* value < 0.05 and a fold change ≥ 1.5 or ≤ -1.5 were defined as the significantly altered compounds in the plots. Compared with the CO_2_ (−) condition, 37, 63, and 78 altered compounds were identified in the early, mid, and late logarithmic phases, respectively; among these compounds, the levels of 24, 39, and 24 compounds increased, and those of 13, 24 and 54 compounds decreased under the CO_2_ (+) condition during the corresponding growth phases. The number of altered metabolites increased towards the late logarithmic phase, suggesting time-dependent changes in the metabolic profile. Under the CO_2_ (+) condition, various metabolic shifts were observed across the early, mid, and late logarithmic phases. All the metabolites identified at each growth phase are listed in Tables S3–S5.

**Figure 2.**
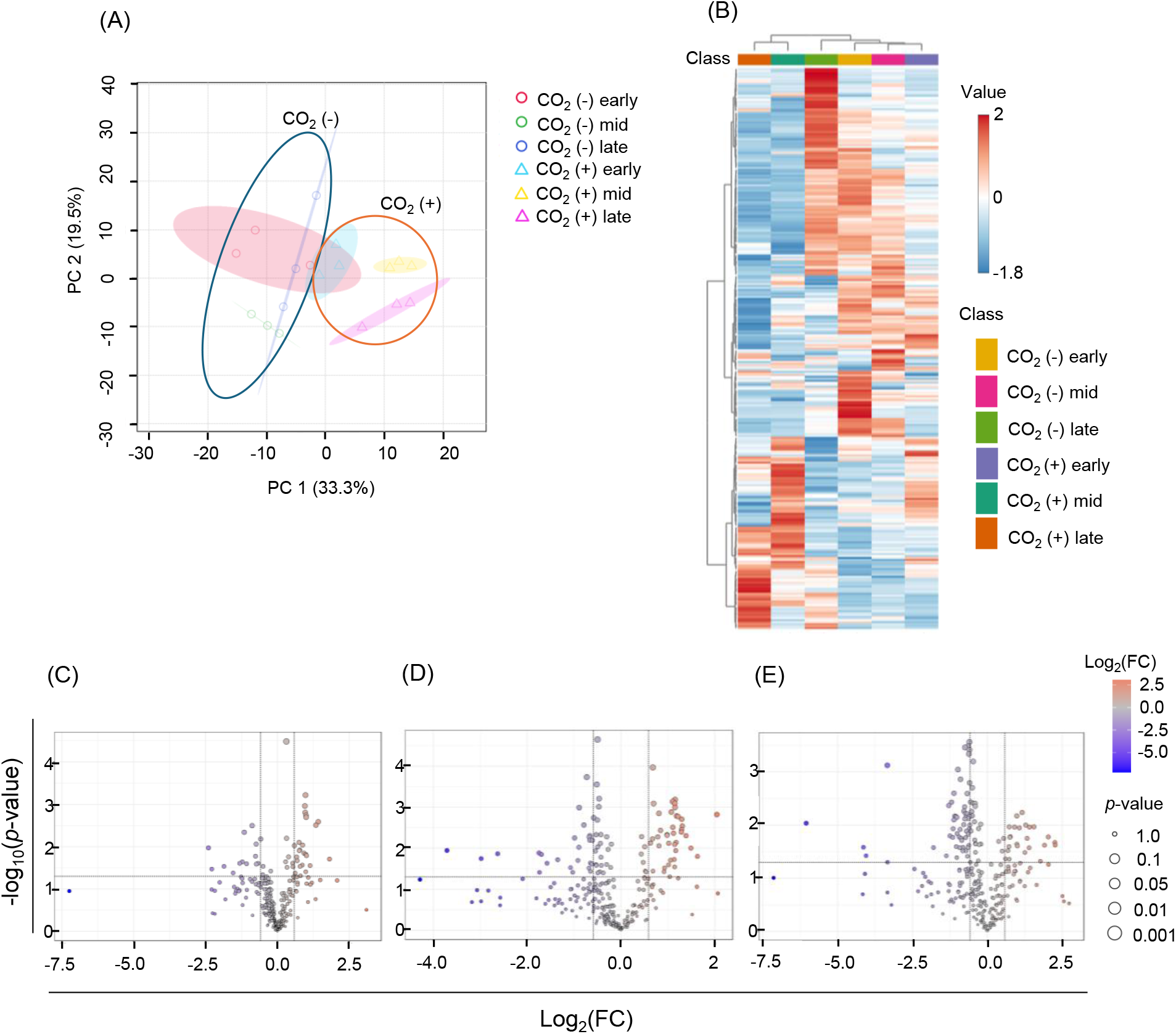
Metabolic profiles of *A. muciniphila* in the presence (+) or absence (−) of CO_2_. (A) Principal component analysis (PCA) plot showing the clustering of metabolic profiles by growth phase and CO_2_ condition. (B) Heatmap showing the results of the hierarchical clustering analysis of significantly altered metabolites. (C)–(E) Volcano plots comparing metabolite abundance between the CO_2_ (+) and CO_2_ (−) groups in the early (C), middle (D), and late (E) logarithmic phases. Significantly altered metabolites were defined as those whose raw *p* value was < 0.05 and whose absolute fold change was ≥ 1.5.

[Figure 2. Metabolic profiles of *A. muciniphila* in the presence (+) or absence (−) of CO_2._]

### Stable ^13^CO_2_ tracing analysis

The results described above revealed that CO_2_ affected the growth and metabolic profile of *A. muciniphila*. However, how *A. muciniphila* utilizes CO_2_ in its metabolism remains unclear. A ^13^C tracing analysis using ^13^CO_2_ was performed to address this issue. Cells grown under ^12^CO_2_ and ^13^CO_2_ conditions at each growth phase were used to calculate the incorporation ratio and number of ^13^C atoms. The ^13^C-incorporated metabolites are shown in Figure 3 and Table S6 and S7. Isotope tracing revealed that ^13^CO_2_-derived ^13^C was extensively incorporated into diverse metabolites through major pathways, including the TCA cycle (acetyl-CoA, malic acid, fumaric acid and succinic acid), de novo synthesis of pyrimidines (UMP, UDP, cytosine, and cytidine 5’-monophosphate), synthesis of TCA-related amino acids (L-aspartate, D-(–)-glutamine, L-pyroglutamic acid, and L-(+)-citrulline), de novo synthesis of purines (adenine, AMP, ADP, ATP, dATP, adenylosuccinic acid, cyclic ADP-ribose, GMP, GDP, and GTP), synthesis of coenzymes (nicotinamide, NAD^+^, NADP^+^, NADPH, CoA, S-adenosylmethionine, and 5‘-S-methyl-5’-thioadenosine and FAD), and synthesis of amino sugars (UDP-GlcNAc and UDP-galactose). Overall, ^13^C incorporation tended to decrease over time from the early to the late exponential growth phase across the ^13^C-labelled metabolites. Furthermore, regarding the ^13^C-incorporated metabolites, the peak areas were compared between the CO_2_ (+) and CO_2_ (−) groups using an untargeted metabolomics dataset (Figure S2). The results showed that not all the metabolites were detected in significantly higher amounts under CO_2_ (+) conditions than under CO_2_ (−) conditions, even when ^13^C was incorporated into these metabolites. Figure 4A and B present a metabolic pathway map inferred from the *A. muciniphila* genome, onto which the results of the untargeted metabolomic analysis and stable ^13^CO_2_ tracing analysis are overlaid.

**Figure 3.**
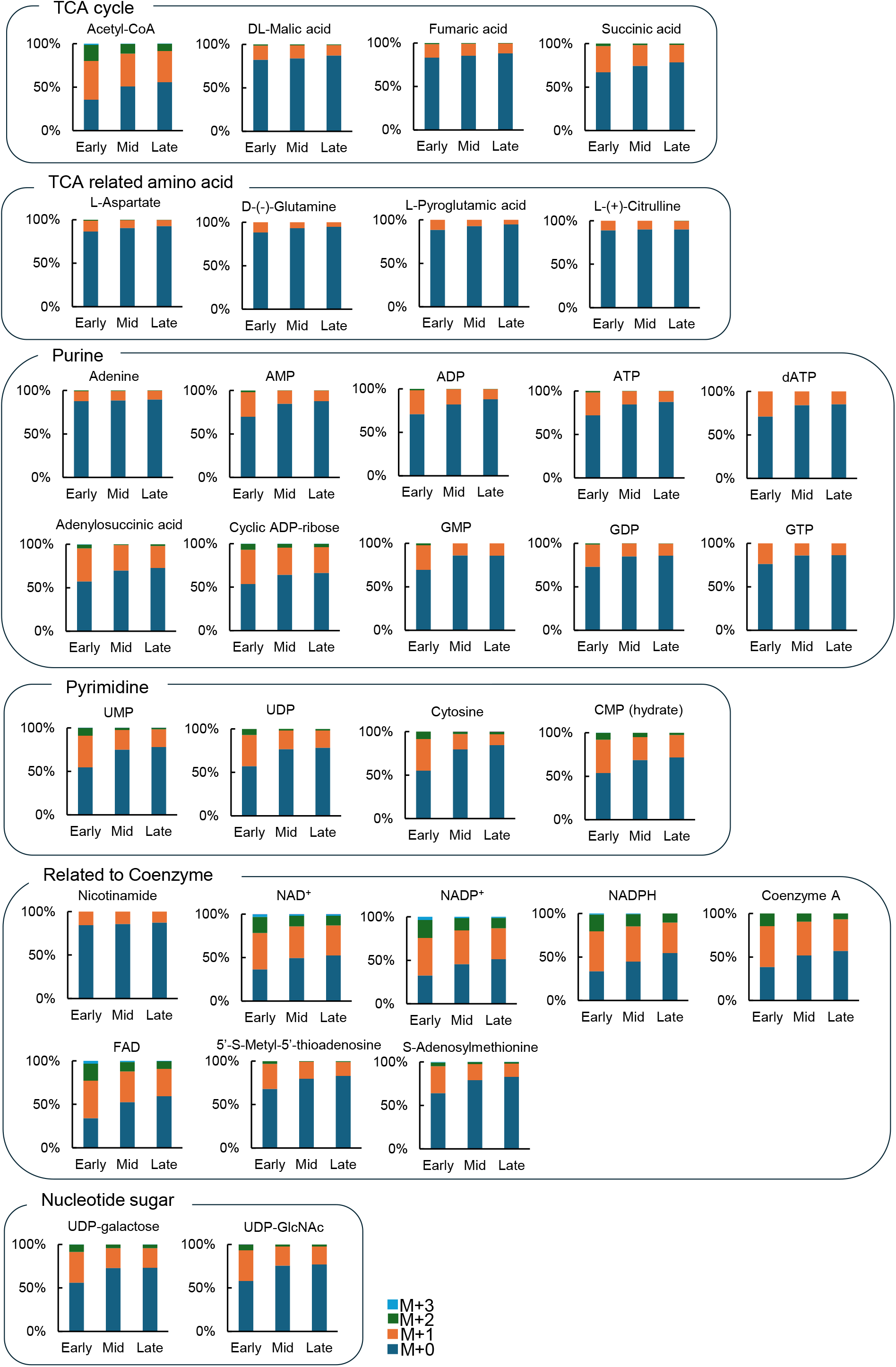
^13^C metabolic tracing analysis using ^13^CO_2_. The incorporation ratio and the number of ^13^C atoms incorporated into each compound are shown in panels categorized by metabolite class. For each bar, the number of incorporated ^13^C atoms is indicated and arranged from bottom to top as M+0 to M+3.

**Figure 4.**
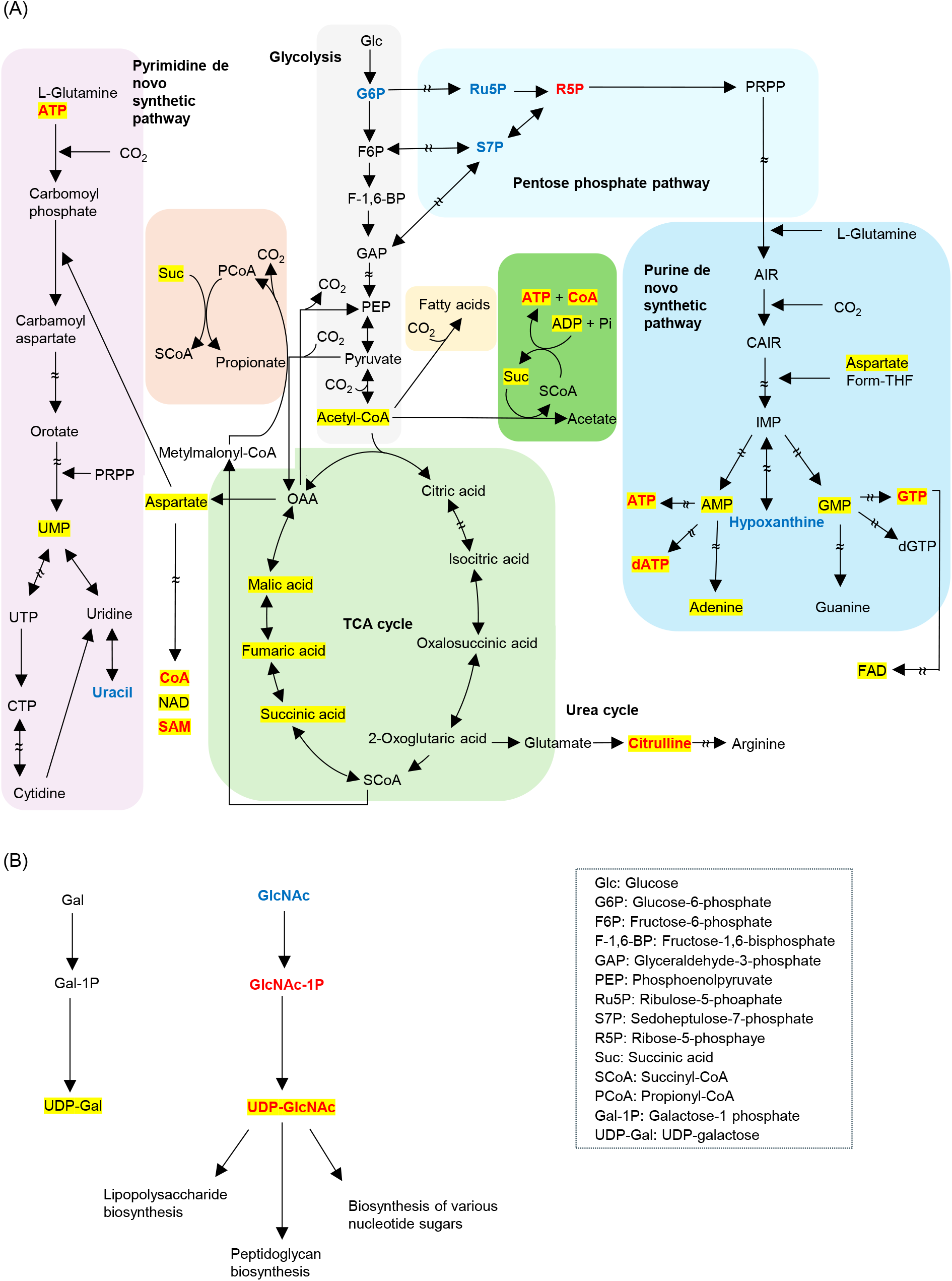
Summary of the results of the metabolomic analysis mapped onto the estimated metabolic pathways in *A. muciniphila*. (A) Central metabolism, nucleic acid metabolism, and organic acid metabolism. (B) Amino sugar nucleotide sugar metabolism. The metabolites shown in blue and red were downregulated and upregulated in the CO_2_ (+) group, respectively (a raw *p* < 0.05, FC ≥ 1.5). Yellow indicates metabolites incorporating ^13^C-derived ^13^CO_2_. These maps were constructed based on the KEGG pathway maps for *A. muciniphila* and a previous report (Ottman *et al*., 2017).

[Figure 3. ^13^C metabolic tracing analysis using ^13^CO_2_.]

[Figure 4. Summary of the results of the metabolomic analysis mapped onto the estimated metabolic pathways in *A. muciniphila*.]

### Measurement of pyruvate carboxylase activity, an anaplerotic reaction associated with carbon fixation

The phosphoenolpyruvate–pyruvate–OAA node in central metabolism is considered an anaplerotic node that replenishes TCA intermediates to direct carbon flux in appropriate directions and fixes CO_2_ via carboxylation (Sauer and Eikmanns, 2005). Recently, pyruvate carboxylase (PCx), an anaplerotic enzyme, was annotated in the *A. muciniphila* genome using ModelSeed (Kim *et al*., 2023). We investigated whether *A. muciniphila* PCx is involved in CO_2_ utilization by measuring PCx activity in CFEs from *A. muciniphila* grown under CO_2_ (−) and CO_2_ (+) conditions. The results revealed that the PCx activity under CO_2_ (+) conditions (204.08 ± 69.32 mU/mg protein) was significantly higher than that under CO_2_ (−) conditions (76.43 ± 23.46 mU/mg protein) (Figure 5).

**Figure 5.**
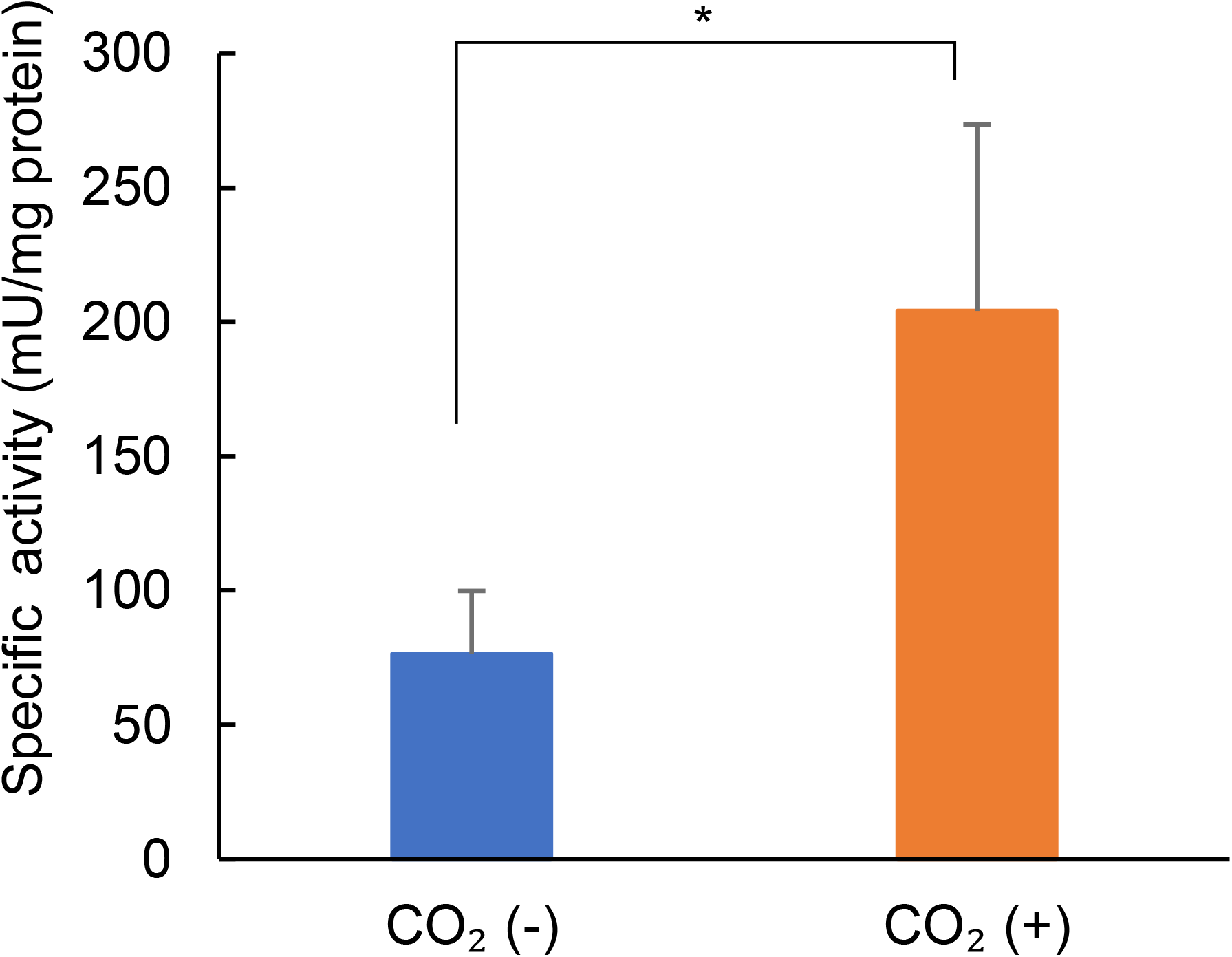
Pyruvate carboxylase (PCx) activity in *A. muciniphila* grown under CO_2_ (−) and (+) conditions. The data are presented as the means ± SDs of four biological replicates, with each measurement performed in technical triplicate. Statistical significance was determined by a two-tailed unpaired Student’s t test (\**p* < 0.05).

[Figure 5. Pyruvate carboxylase (PCx) activity in *A. muciniphila* grown under CO_2_ (−) and (+) conditions.]

### Oxaloacetate (OAA) supplementation in *A. muciniphila* cultures

The results thus far indicate both the incorporation of CO_2_-derived carbon into L-aspartate and malic acid (which are interconverted with OAA) (Figure 3) and higher PCx activity in *A. muciniphila* grown under CO_2_ (+) conditions (Figure 5). On the basis of these results, we hypothesized that the production of OAA via the anaplerotic PCx reaction contributes to the increased growth of *A. muciniphila*. We investigated this possibility by performing additional experiments in which the cultures were supplemented with 0–10 mM OAA instead of CO_2_. The results showed that supplementation with OAA increased the growth of *A. muciniphila* in a dose-dependent manner, similar to that observed with CO_2_ supplementation (Figure 6A and B).

**Figure 6.**
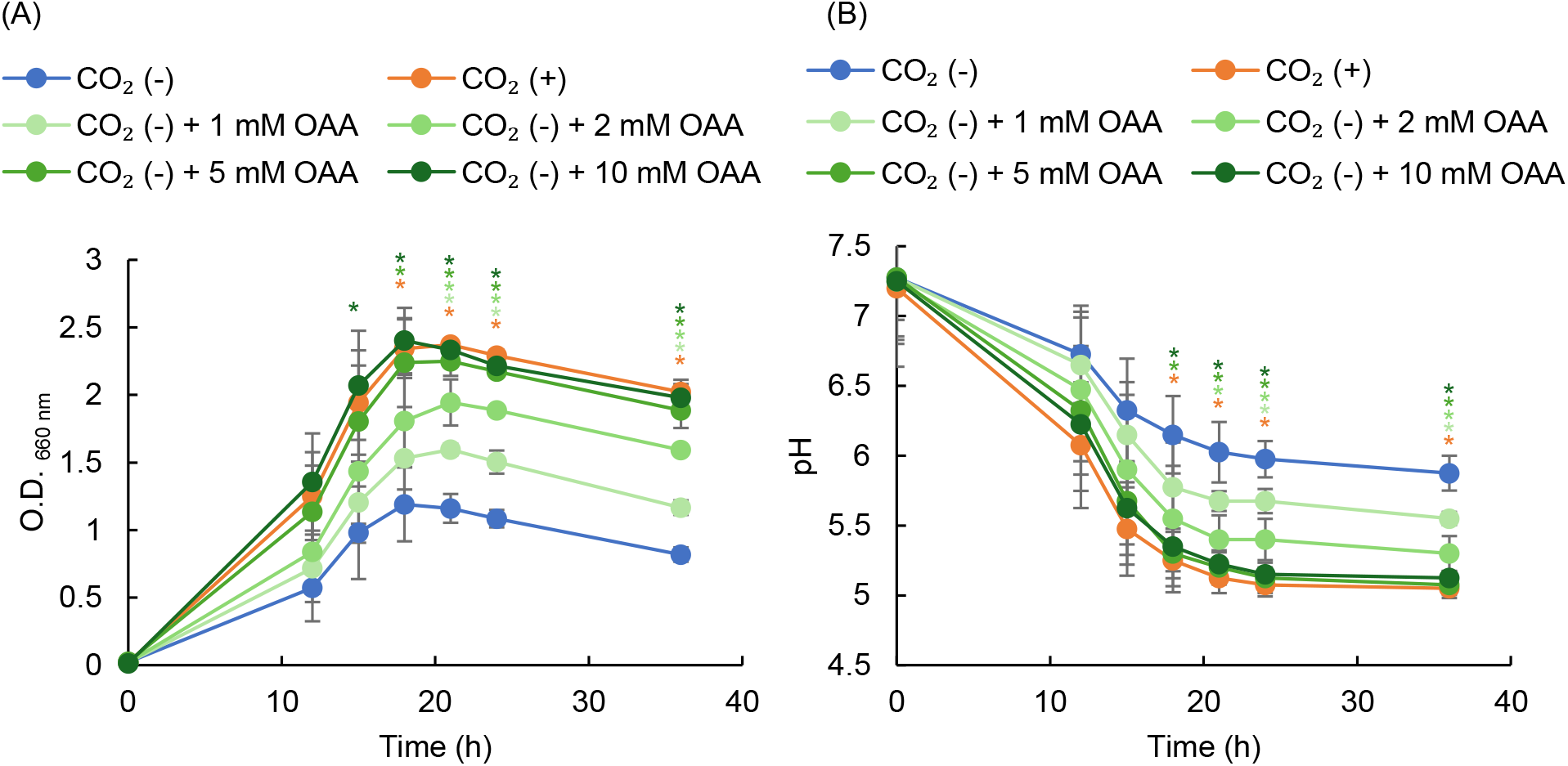
Effect of OAA on the growth of *A. muciniphila*. (A) Growth of *A. muciniphila* under CO_2_ (−) conditions in medium supplemented with OAA (0, 1, 2, 5, and 10 mM) and under CO_2_ (+) condition. (B) pH measured during culture. The data are presented as the means ± SDs of three biological replicates. Statistical significance was determined by Tukey’s HSD test (\**p* < 0.05, vs. 0% CO_2_).

[Figure 6. Effect of OAA on the growth of *A. muciniphila*.]

## 4. Discussion

The gut microbiota is related to host health, and CO_2_ has been empirically utilized to culture such anaerobes from the gut. However, its detailed effects on these anaerobes remain unclear, although intestinal CO_2_ levels vary among individuals depending on the host’s diet and physiological status. In this study, we investigated the effect of CO_2_ on a beneficial microbe in the gut, *A. muciniphila*. We demonstrated that the presence of CO_2_ increased its growth, induced a metabolic shift and promoted the active incorporation of CO_2_ into its metabolism. In particular, this CO_2_ incorporation suggests that the turnover of anaplerotic TCA cycle intermediates and coenzymes is enhanced, thereby enabling smooth metabolic flux to support cell growth.

In the presence of CO_2_, *A. muciniphila* exhibited increased growth and a sharp decrease in pH over time compared with the absence of CO_2_ in GAM and CP media (Figure 1A and B, Figure S1A and B). These results are similar to a previous report showing that CO_2_ supplementation in Brain Heart Infusion-Mucin (BHIM) medium promoted the growth of *A. muciniphila* (Wang *et al*., 2025). Thus, these results suggest that CO_2_ is required for the optimal growth of *A. muciniphila* in diverse culture media.

The TCA cycle serves as a central hub of both anabolic and catabolic processes, and its intermediates are withdrawn through cataplerotic reactions and used as precursors for the biosynthesis of macromolecules, including nucleic acids, proteins, polysaccharides, lipids, and coenzymes. Therefore, maintenance of TCA cycle function requires a continuous supply of acetyl-CoA and replenishment of TCA cycle intermediates through anaplerotic reactions, particularly those supplying OAA (Arnold and Finley, 2023). With respect to the TCA cycle, the incorporation of ^13^C into acetyl-CoA, malic acid, fumaric acid, and succinic acid was observed (Figure 3). Notably, a high level of ^13^C incorporation in coenzymes was detected. Since the main end products were acetic acid and succinic acid, while propionate was minor (data not shown), it was considered possible that succinic acid served as the end product without progressing to propionate in order to consume excess reducing equivalents such as coenzymes including FADH_2_ and NADH generated by CO_2_ utilization. Furthermore, *A. muciniphila* grown under the CO_2_ (+) condition exhibited increased PCx activity, which is involved in a major anaplerotic reaction of the TCA cycle to generate OAA from pyruvate and CO_2_ (Figure 5). This CO_2_-derived carbon is incorporated into the -C4 (−COO^-^) of OAA (Utter and Keech, 1963). In addition, supplementation with OAA stimulated the growth of *A. muciniphila* under the CO_2_ (−) condition, similar to that observed under the CO_2_ (+) condition. Although OAA was not detected in our analysis, because it is unstable and rapidly underwent nonenzymatic decomposition while metabolite extraction, its downstream aspartic acid and compounds in the reductive branch of the TCA cycle (such as malic acid, fumaric acid, and succinic acid) exhibited ^13^C incorporation (Figure 3). These results suggest that *A. muciniphila* utilizes CO_2_ to generate OAA via anaplerosis and to increase TCA cycle flux.

Furthermore, the incorporation of ^13^CO_2_-derived carbon into compounds within purine and pyrimidine metabolic pathways, as well as adenine-related coenzymes, was observed (Figure 3). Aspartate, which is produced from OAA, also exhibited ^13^C incorporation and might be involved in the synthesis of these compounds because aspartic acid can be incorporated into phosphoribosylaminoimidazolesuccinocarboxamide (SAICAR), a precursor of purines. However, the carbon incorporated into aspartic acid from OAA is not retained in the final purine product because only an amino group of aspartate is incorporated into the purine ring, while its carbon skeleton is released (e.g., as fumaric acid) (Levenberg *et al*., 1956) (Miller *et al*., 1959). Thus, direct CO_2_ fixation, such as 5-aminoimidazole ribonucleotide (AIR) carboxylation by AIR carboxylase, may explain this ^13^C incorporation in purine compounds (Figure 4A). Adenine produced via this purine biosynthesis pathway and aspartate from OAA also contribute to coenzyme biosynthesis, which might result in the observed incorporation of ^13^C into coenzymes. On the other hand, the carbon skeleton in aspartate (C4–C6) (Kishore *et al*., 2005) is likely retained during pyrimidine biosynthesis in *A. muciniphila*. The incorporation of OAA-derived aspartate accounts for one CO_2_-derived ^13^C atom in pyrimidine compounds. Since some pyrimidine compounds incorporated two ^13^C atoms, other CO_2_ fixation pathways might be involved in this ^13^C labelling of pyrimidine compounds, such as the carbamoyl phosphate synthetase (CPS) reaction, which introduces CO_2_ to carbamoyl phosphate. These pyrimidines are also involved in nucleotide sugar metabolism; indeed, UDP-GlcNAc and UDP-galactose incorporated ^13^C, suggesting that CO_2_-derived carbon flows into the glycosylation pathway. These results suggest that under CO_2_ (+) conditions, *A. muciniphila* utilizes CO_2_ for the production of various metabolites via the de novo purine/pyrimidine synthesis pathways and coenzyme metabolism, which are essential for the maintenance of multiple metabolic pathways.

*A. muciniphila* was able to grow in GAM medium even in the absence of CO_2_, reaching approximately half the growth level observed in the presence of CO_2_. Since, compared with CP medium, GAM medium is nutritionally rich and contain large amounts of extracts that supply nucleic acids, amino acids, and lipids, they might enable *A. muciniphila* to utilize these components directly without relying on CO_2_-dependent biosynthesis, thereby achieving rapid growth even under the CO_2_ (−) condition. In contrast, in CP medium, growth was delayed under the CO_2_ (−) condition. This result might be because *A. muciniphila* utilized self-produced CO_2_ via its own metabolism under nutrient-poor conditions. Recently, *A. muciniphila* was reported to exacerbate food allergy in mice under dietary fibre-deficient conditions (Parrish *et al*., 2023). In addition, *A. muciniphila* has been shown to reduce the pathogenicity of *C. rodentium* in the presence of dietary fibre (Wolter *et al*., 2024). Since dietary fibre intake can increase CO_2_ levels in the human gut (Modesto *et al*., 2022), these findings suggest that the physiological status of *A. muciniphila* and its effect on host health may vary depending on diet-induced changes in intestinal CO_2_ levels within the gut ecosystem.

In conclusion, our results suggest that *A. muciniphila* utilizes CO_2_ for OAA production via the PCx-catalysed anaplerotic reaction in the TCA cycle and other pathways involving CO_2_ incorporation. OAA and its metabolite aspartate serve as precursors for the de novo purine/pyrimidine synthesis pathway and as carbon sources for coenzyme synthesis, both of which are essential for driving and sustaining multiple metabolic pathways, supplying metabolic intermediates continuously, and maintaining metabolic flux. Through these metabolic pathways, CO_2_ may promote the robust growth of *A. muciniphila*. These mechanisms may contribute to the effects of *A. muciniphila* on host interactions, and similar phenomena could occur in other gut microbiota. Further studies are needed to achieve a comprehensive understanding of the effects of CO_2_.

## Acknowledgements and funding

This study was partly funded by the Tokyo University of Agriculture Doctoral Research Support Program. AI was used solely for English-language editing. All scientific content was reviewed and verified by the authors.

## CRediT authorship contribution statement

Conceptualization, MS, SO and TS; methodology, MS, SO and TS; validation, MS, SO, RO and JY; formal analysis, MS; resources, MS and TS; writing—original draft preparation, MS; writing—review and editing, TS; visualization, MS; supervision, TS; project administration, TS; funding acquisition, MS; investigation, MS, SO, RO and JY. All the authors have read and agreed to the published version of the manuscript.

