## Supplementary FIgure for "Effects of CO_2_ on the growth and metabolism of *Akkermansia muciniphila*"

**Preparation of culture medium**

CP medium were also used with minor modifications to further investigate the effects of $CO_2$. CP medium was prepared based on the composition described previously (Derrien *et al.*, 2004). Briefly, 1 liter of medium contained 0.4 g $KH_2PO_4$, 0.53 g $Na_2HPO_4$ 12 ($H_2O$), 0.3 g $NH_4Cl$, 0.3 g NaCl, 0.1 g $MgCl_2$ 6 ($H_2O$), 0.11 g $CaCl_2$ 2 ($H_2O$), 1 mL an acidic trace element solution, and 1 mL alkaline trace element solution, as described previously, but lacked $NiCl_2$ and $Na_2SeO_3$. Before inoculation, the medium was supplemented with 25 mM GlcNAc, 0.1 g/L glucose, 0.05% L-cysteine HCl, 11 g/L $CaCl_2$ $2H_2O$, 1 ml/L standard vitamin solution, 4 g/L L-threonine, and 47.6 mM MOPS (pH 8.0). MOPS was used as a substitute for $NaHCO_3$. *A. muciniphila* was incubated in CP medium at 37 ° C with orbital shaking at 45 rpm in the presence of 0, 1, 3, 5, 10, and 20% $CO_2$.

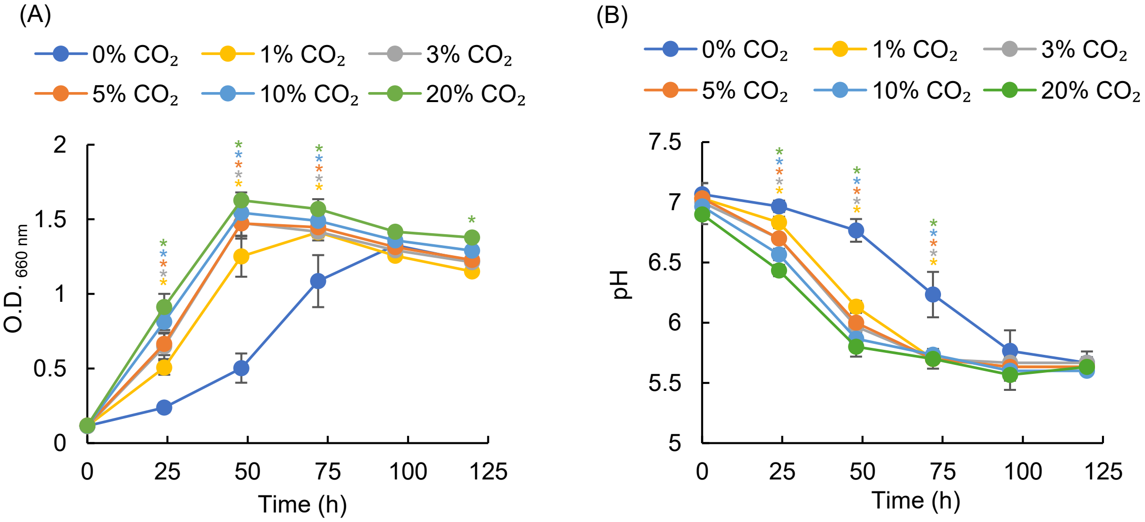


Figure S1. Effect of $CO_2$ on the growth *A. muciniphila* in CP medium.
(A) Growth of *A. muciniphila* and changes in pH (B) in the presence of 0%, 1%, 3%, 5%, 10% and 20% $CO_2$ in CP medium. (A) and (B) are presented as means ± SDs of three biological replicates. Statistical significance was determined by Tukey's HSD test (*$p$ < 0.05, vs. 0% $CO_2$).

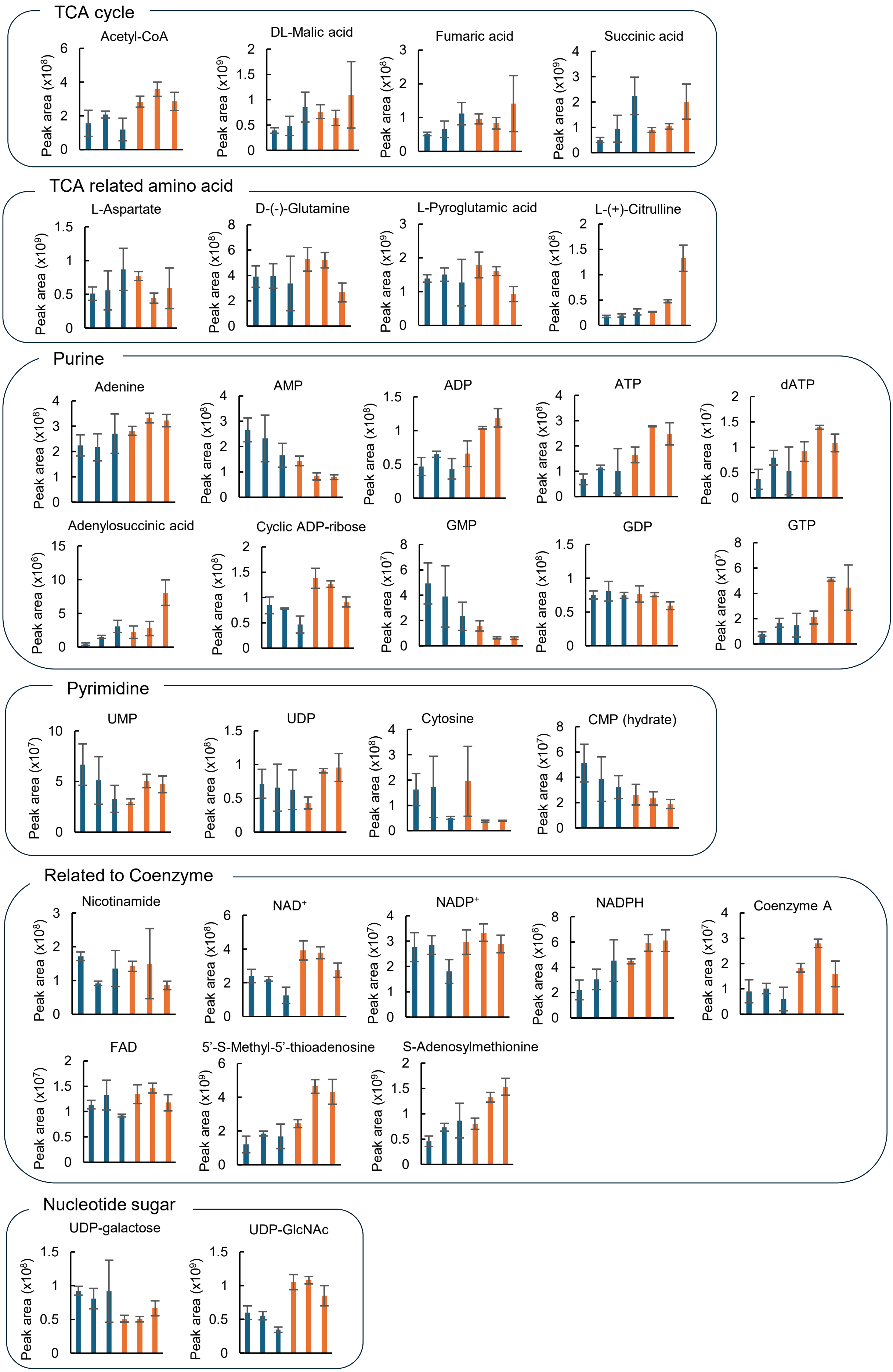


Figure S2. Stable isotope tracing analysis of *A. muciniphila* using $^{13}CO_2$.
Bar graphs representing the relative abundance of metabolites in which incorporation of $^{13}C$ was observed, as shown in Figure 3. The data are presented as means ± SDs of three biological replicates. The graphs are arranged from left to right as follows: the early, mid, and late logarithmic phases of $CO_2$ (-), followed by the early, mid, and late logarithmic phases of $CO_2$ (+).
